# EasyCircR: Detection and reconstruction of circular RNAs post-transcriptional regulatory interaction networks

**DOI:** 10.1101/2023.12.12.571275

**Authors:** Antonino Aparo, Simone Avesani, Luca Parmigiani, Sara Napoli, Francesco Bertoni, Vincenzo Bonnici, Luciano Cascione, Rosalba Giugno

**Affiliations:** Department of Computer Science, University of Verona, Verona, 37134, Italy; Faculty of Technology and Center for Biotechnology (CeBiTec), Bielefeld University, Bielefeld, 33615, Germany; Bielefeld Institute for Bioinformatics Infrastructure (BIBI), Bielefeld University, Bielefeld, 33615, Germany; Graduate School “Digital Infrastructure for the Life Sciences” (DILS), Bielefeld University, Bielefeld, 33615, Germany; Institute of Oncology Research, Bellinzona, 6500, Switzerland; Oncology Institute of Southern Switzerland (IOSI), Bellinzona, 6500, Switzerland; Department of Mathematical, Physical and Computer Sciences, University of Parma, Parma, 43124, Italy; Swiss Institute of Bioinformatics (SIB), Lausanne, 1015, Switzerland

## Abstract

Circular RNAs (circRNAs) are regulatory RNAs that play a crucial role in various biological activities and have been identified as potential biomarkers for the development of neurological disorders and cancer. CircRNAs have emerged as significant regulators of gene expression through different mechanisms, including regulation of transcription and splicing, modulation of translation, and post-translational modification. Additionally, some circRNAs function as microRNA (miRNA) sponges in the cytoplasm, enhancing the expression of target genes by suppressing miRNA activity. Although existing pipelines reconstruct circRNAs, identify miRNAs sponged by them, retrieve cascade-regulated mRNAs, and represent the regulatory interactions as complex circRNA-miRNA-mRNA networks, none of the state-of-the-art approaches can discriminate the biological level at which the mRNAs involved in the interactions are regulated.

EasyCircR is a novel R package that combines circRNA detection and reconstruction with post-transcriptional gene expression analysis (exon-intron split analysis) and miRNA response element prediction. The package enables estimation and visualization of circRNA-miRNA-mRNA interactions through an intuitive Shiny application, taking advantage of the post-transcriptional regulatory nature of circRNAs and excluding unrealistic regulatory outcomes. EasyCircR source code, Docker container and user guide are available at: https://github.com/InfOmics/EasyCircR

## 1. Introduction

Circular RNAs (circRNAs) are a type of non-coding single-stranded RNAs that play a vital role in a wide range of biological processes, including cell proliferation, differentiation, and development. Additionally, they have been linked to cancer initiation and progression, and their resistance to endonucleases makes them excellent biomarker candidates. Unlike linear RNAs, long non-coding RNAs, and microRNAs (miRNAs), circRNAs have covalently closed loop structures without free 3’ poly(A) tails or 5’ caps. They contain multiple miRNA response elements (MREs) that enable them to bind miRNAs and act as competitive endogenous RNAs. By sponging miRNAs, circRNAs can reduce the inhibition of miRNA-targeted genes, consequently modulating their expression level [1]. Studies have shown that the circRNAmiRNA-mRNA interaction network is dysregulated in the pathogenesis of many diseases, including cancer. Therefore, after circRNA detection [2], it is biologically important to infer the circRNA-miRNA-mRNA network by predicting the sponged miRNAs and the putatively regulated target genes [3, 4].

Although several pipelines for circRNA-miRNA-mRNA interaction estimation already exist [5, 6], only a few of them allow for the inference of different circRNA regulatory effects between two sample conditions, and none of the released pipelines take advantage of the post-transcriptional regulatory nature of the circRNAs, which can lead to potentially incorrect interactions in the results. In this context, we developed EasyCircR, an R package that allows users to easily perform the entire circRNA regulation analysis and visualize interaction results through an intuitive graphical interface. The tool reconstructs full-length circRNAs from FASTQ files collected from samples of several groups (e.g., Control vs Tumor, Treated vs Untreated), retrieves the genes that are modulated between the conditions of interest, and predicts their binding miRNAs. To filter out potential unrealistic regulatory interactions, only the post-transcriptionally regulated genes are kept in the analysis. To search for them, EasyCircR applies an exon-intron split analysis [7] that detects variations in transcriptional and post-transcriptional regulation by estimating differences in exonic and intronic changes across different conditions. Finally, EasyCircR provides an interactive R Shiny application where regulatory interactions can be examined individually, filtered based on specific criteria, and exported in textual format for additional research. EasyCircR was tested on an RNA-seq dataset collected from diffuse large B cell lymphoma (DLBCL) cell line [8] exposed to Bimiralisib, an inhibitor with preclinical and early clinical anti-lymphoma activity [9]. Interestingly, the results revealed, in the treated samples, the down-regulation of a tumor promoter circRNA and the up-regulation of circRNAs (Table 1) which induce the positive post-transcriptional regulation of tumour suppressor genes capable of inhibiting the growth and diffusion of tumour cells.

**Table 1:** Table showing the significant differential expressed circRNAs. For each circRNA are shown the Ensembl ID and symbol of the gene where the circRNA locates, the statistics results of circRNAs differential expression analysis including log fold change, average expression, p-value and adjusted p-value, the validated name of the circRNA retrieved form circBank and the number of harboured miRNAs.

| circRNA (BSJ ID) | Ensembl ID | Gene Name | logFC | AveExpr | p-value | adj.p-value | validated | #miRNA |
| --- | --- | --- | --- | --- | --- | --- | --- | --- |
| 1:247155566—247159813:-:1 | ENSG00000196418 | ZNF124 | -5.121 | 11.153 | 2.459347e-12 | 1.337e-09 | hsa_circZNF124.005 | 201 |
| 2:61522611—61533903:- | ENSG00000082898 | XPO1 | -4.050 | 11.013 | 6.593850e-07 | 1.793e-04 | hsa_circXPO1.001 | 170 |
| 3:196391813—196403019:- | ENSG00000163960 | UBXN7 | 3.0526 | 12.879 | 1.807034e-06 | 2.579e-04 | hsa_circUBXN7.003 | 129 |
| 7:116092137—116112038:-:3 | ENSG00000105967 | TFEC | 3.379 | 11.069 | 3.429041e-06 | 2.579e-04 | - | 220 |
| 7:116092137—116112038:-:7 | ENSG00000105967 | TFEC | 3.379 | 11.069 | 3.429041e-06 | 2.579e-04 | - | 235 |
| 7:116092137—116112038:-:11 | ENSG00000105967 | TFEC | 3.379 | 11.069 | 3.429041e-06 | 2.579e-04 | - | 296 |
| 7:116092137—116112038:-:15 | ENSG00000105967 | TFEC | 3.379 | 11.069 | 3.429041e-06 | 2.579e-04 | - | 311 |
| 18:9182382—9221999:+ | ENSG00000265257 | - | 4.566 | 11.429 | 3.793828e-06 | 2.579e-04 | hsa_circANKRD12.009 | 347 |
| 12:116230533—116237705:-:13 | ENSG00000265257 | MED13L | 3.251 | 10.809 | 3.062081e-05 | 1.388e-03 | hsa_circMED13L.005 | 229 |
| 12:116230533—116237705:-:16 | ENSG00000265257 | MED13L | 3.251 | 10.809 | 3.062081e-05 | 1.388e-03 | hsa_circMED13L.005 | 269 |

## 2. Methods

EasyCircR is a powerful tool that detects circRNA-miRNA-mRNA interactions from RNA-seq data. It offers various options for exploring reconstructed circRNA structures and exporting filtered results for further analysis. EasyCircR performs four key steps: identifying differentially expressed (DE) circRNAs, predicting circRNA-miRNA interactions, analyzing post-transcriptional regulation, integrating and visualizing data (see Figure 1).

**Figure 1:**
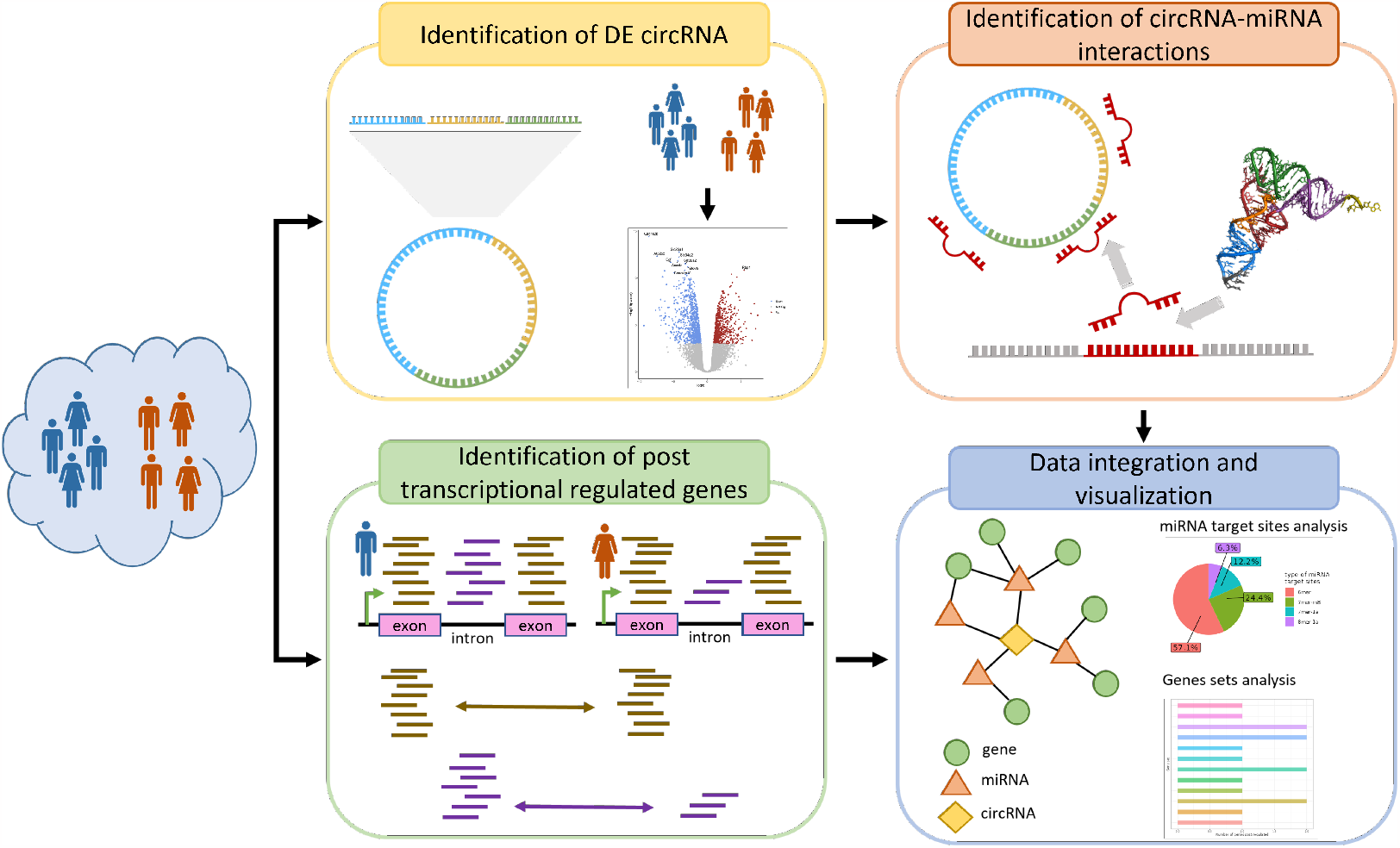
A graphical overview of EasyCircR pipeline: starting from RNA-seq data collected from two samples groups, the package quantifies the circularRNAs, identifies those differentially expressed between the two samples conditions and, for each of them, detects the binding microRNAs. To conclude, EasyCircR infers, starting from the sequencing data, the post-transcriptional regulated genes and constructs putatative circulaRNA-microRNA-gene regulative interactions that could be explored and visualized through an intuitive Shiny app.

Starting from RNA-seq’s FASTQ, in the first step, it begins by reconstructing full-length circRNAs using CIRI-full [10]. CIRI-full utilizes the reverse overlap (RO) and back-splice junction (BSJ) features to identify low-abundance circRNAs. CIRI-full relies on CIRI2 [11] for circRNA detection [12] and CIRI-AS [13] to detect cirexon (circRNA’s exon) and alternative splicing events in circRNAs. Since CIRI-AS cannot process sequencing reads with different lengths, EasyCircR accepts reads having all the same length or trims reads using the Trimmomatic tool [14] according to the user’s desired length passed as a parameter to the package. In order to preserve the highest number of reads, EasyCircR provides a visualization function that allows exploring the distribution of read lengths before trimming. EasyCircR runs CIRI-vis [15] to visualize the alignments of BSJ and RO merged reads and the reconstructed circRNA sequences. After circRNA detection, EasyCircR computes circRNA differential expression analysis (DE) by using limma+voom [16] R packages. Given CIRI-full ability to detect circRNAs at the isoform level, DE is performed considering alternative splicing of transcripts. Additionally, EasyCircR provides the functionality to map significant differentially expressed circRNAs to their host gene annotation [17] supplied by the circBank database [18], which stores over 140,000 human-annotated circRNAs. In the second step, EasyCircR computes the circRNA-miRNA interactions by using TargetScan [19], which predicts miRNA response elements (MREs) from the entire circRNA sequences. As the third step, EasyCircR identifies the post-transcriptionally regulated genes. It uses the EISA algorithm of eisaR package [7], which quantifies genes starting from FASTQ input files and sorts them based on differences in exonic and intronic read counts between conditions. Only statistically significant genes (FDR ¡ 0.05) are kept as potential circRNA target genes. Finally, circRNA-miRNA-mRNA interaction networks are constructed by retrieving the miRNAs targeting the genes returned during the third step using multiMiR [20]. Interactions are visualized through a Shiny app. The app includes multiple types of feature filters together with the possibility of investigating reconstructed circRNA structures and exporting filtered results in textual format for further analysis.

## 3. Using EasyCircR Shiny App

As outlined in the methodology section, upon the completion of computational analysis through EasyCircR, the package provides a Shiny web application for visualizing the outcomes. This application comprises three tables (see Figure 2) containing comprehensive listings of identified circRNAs, miRNAs, and mRNAs resulting from the analysis.

**Figure 2:**
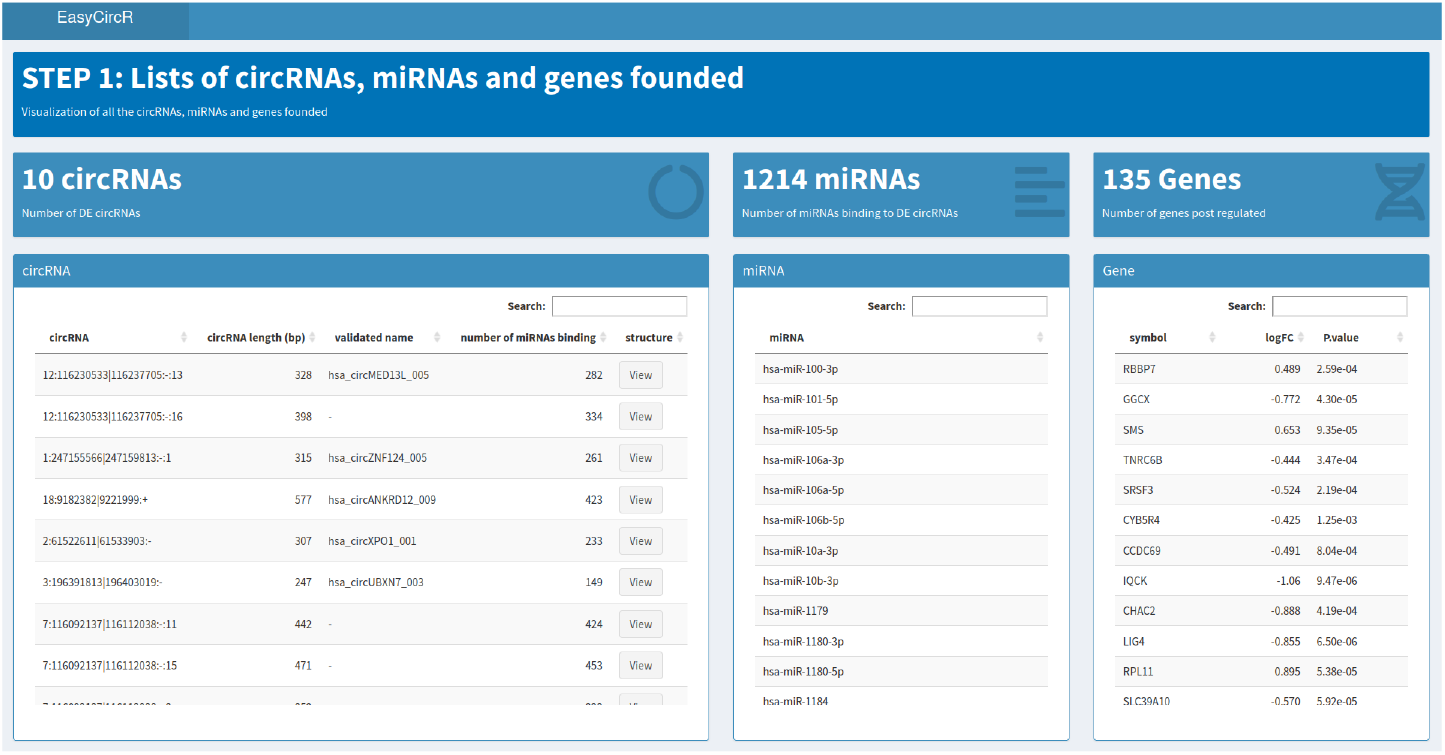
EasyCircR Shiny app showing the complete lists of circRNAs, miRNAs and mRNAs identified in the analysis.

When a specific entry is chosen from any of these tables (such as a gene, miRNA, or circRNA), only the constituent components of the chosen entry regulatory interactions are exhibited (see Figure 3).

**Figure 3:**
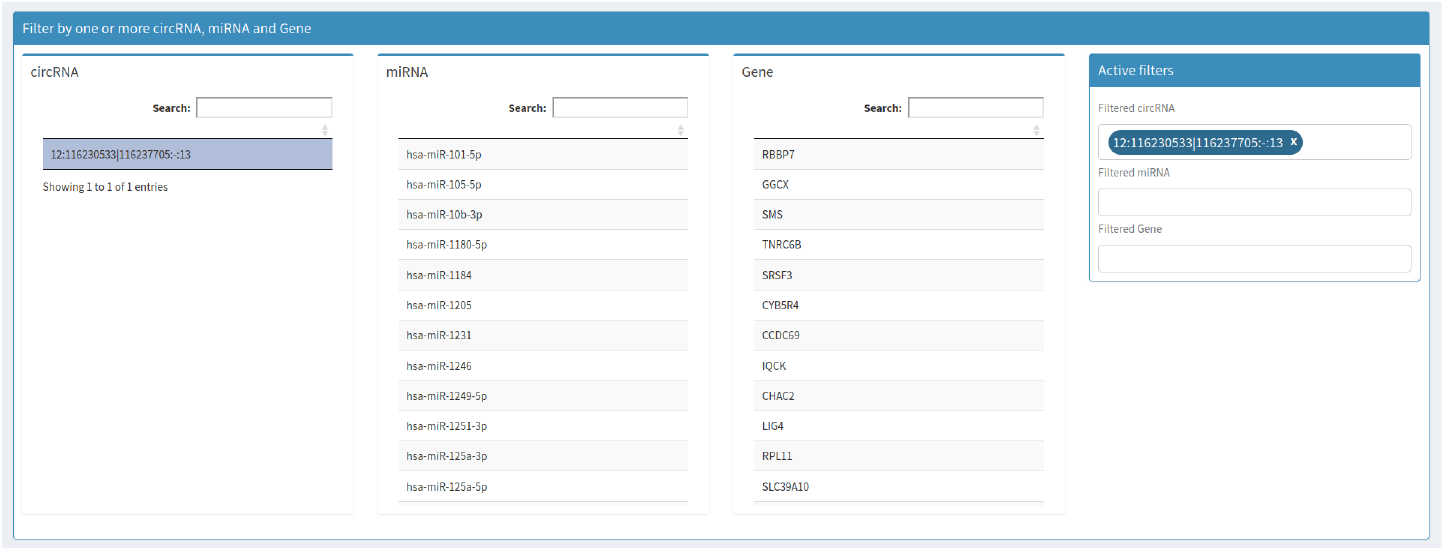
Filtered results. Tables store the elements composing the selected entry interactions.

Furthermore, by activating the “View” button, the structure of the selected circRNA provided by CIRI tool [10] is shown alongside the regions wherein miRNAs are bound (see Figure 4).

**Figure 4:**
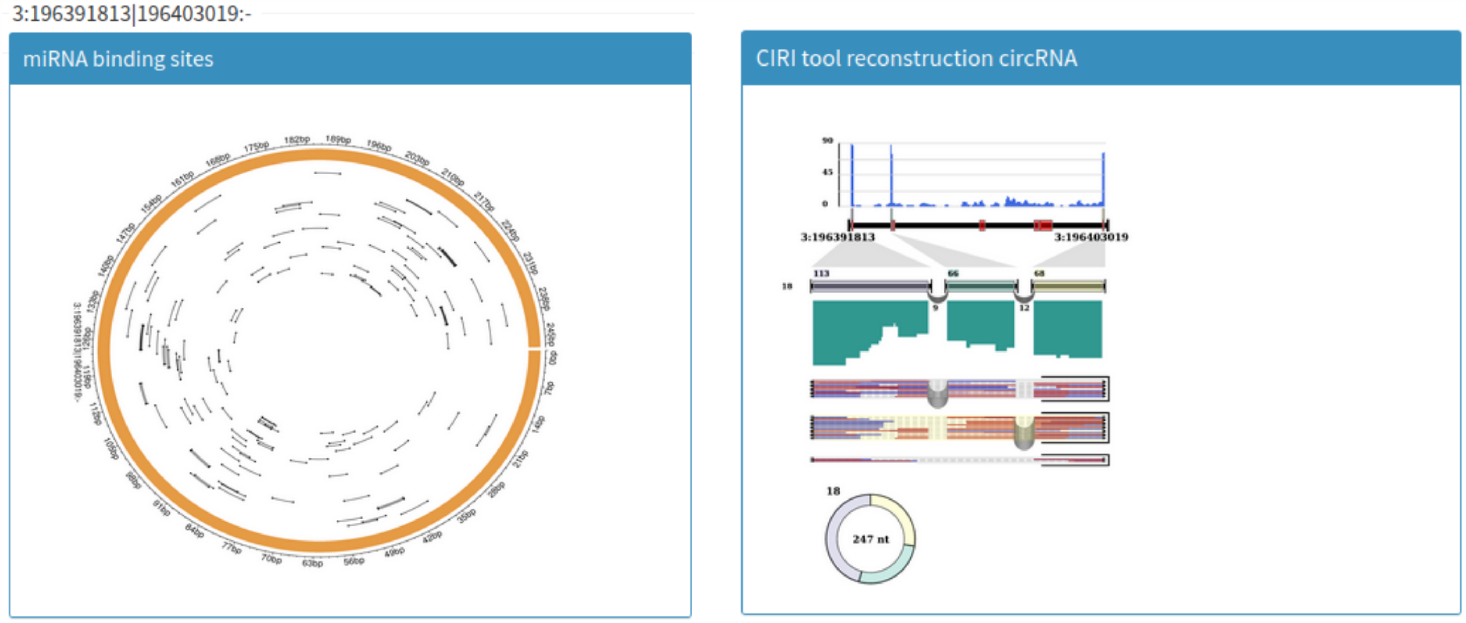
The pop-up page showing the selected circRNA structure and its miRNAs binding sites.

As one scrolls down the web application interface, a filter section becomes accessible (see Figure 5). This filter mechanism permits interactions to be refined using designated criteria like the source database, the circRNA chromosome, initiation or termination positions of the circRNAs, and various others.

**Figure 5:**
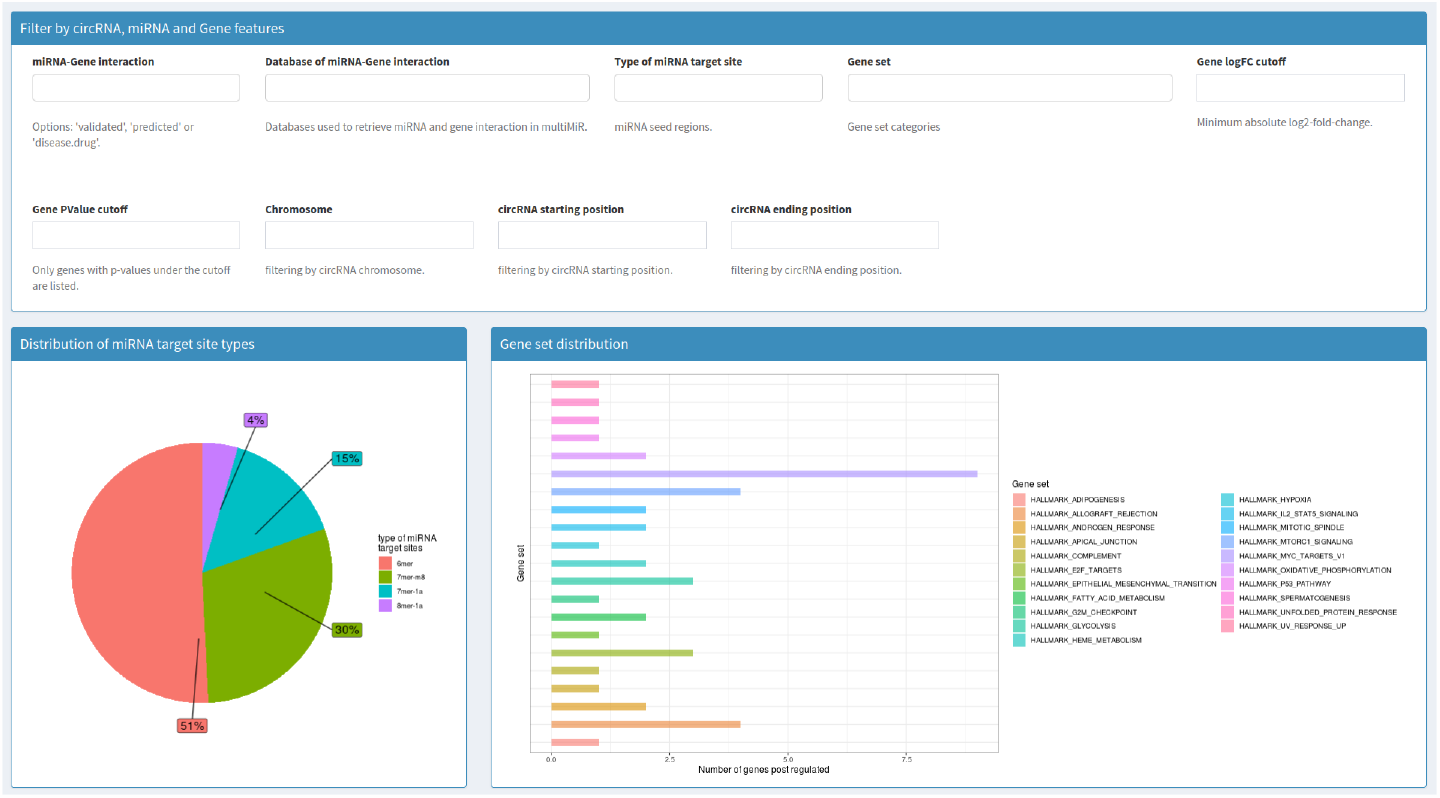
A filter section to restrict the returned results and distributions plots showing miRNAs target sites type and gene sets identified during the analysis.

In tandem with the filter options, the application includes graphical representations illustrating the distributions of miRNA target site types and gene sets.

Positioned at the lower part of the web application page (see in Figure 6), an exhaustive array of potential relationships is cataloged within a navigable table. This table is dynamic, adapting in real-time based on the applied filters. Furthermore, users can export the table in diverse formats (csv, xlsx), facilitating subsequent in-depth analyses.

**Figure 6:**
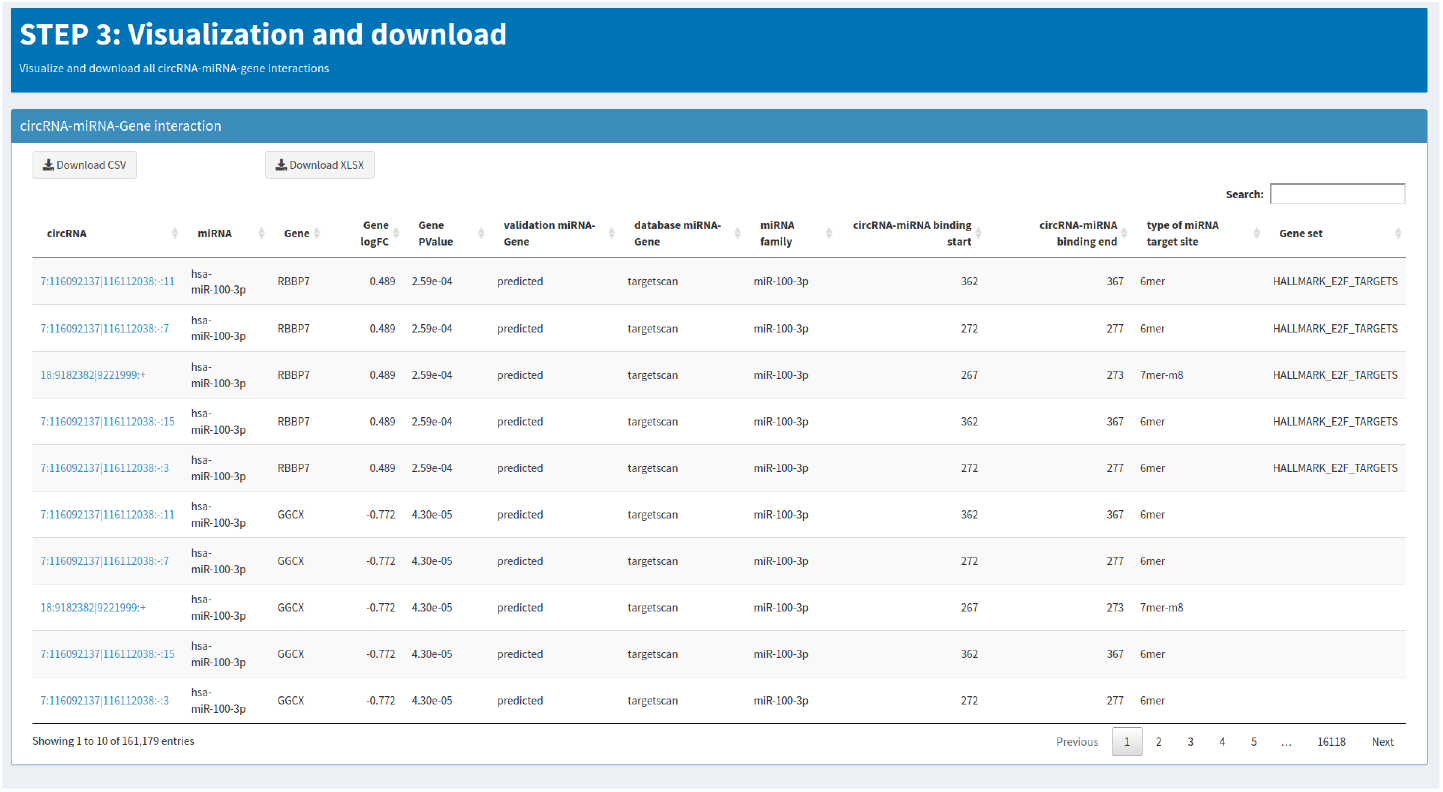
The table storing all the possible relations retrieved during the analysis and downloadable in different formats including csv and xlsx.

## 4. Results

To demonstrate the usefulness of EasyCircR, we analyzed RNA-seq data from DLBCL cell lines [8] treated with Bimiralisib. Our analysis confirmed the post-transcriptional and anti-lymphoma regulatory role of the drug via circRNA-miRNA-mRNA interactions. By comparing treated with untreated samples of the TMD8 cell line, EasyCircR identified 10 significantly differentially expressed circRNAs that potentially sponge 1214 miRNAs and regulate 135 genes at the post-transcriptional level. The analysis revealed circRNAmiRNA-mRNA interactions that support the biological results discussed in [8]. In the treated samples, EasyCircR highlighted the down-regulation of hsa circXPO1 001 circRNA, already cited in the literature as tumor growth promoter [21], and the up-regulation of hsa circUBXN7 003 and four circRNAs on chromosome 7 with binding sites for miR-429 and miR-148b-3p (Table 1). These two miRNAs are already known as biomarkers for cancer [22] and promote cancer cell proliferation and migration. Moreover, miR-148b3p is also known for its ability to target tumor-suppressor genes negatively regulating the PI3K/Akt pathway [23] and promoting tumor growth. Finally, we explored the post-transcriptional regulated genes that compose the circRNA-miRNA-mRNA interaction network, discovering some involved in the Cellular response to starvation Reactome [24] pathway and the regulation of translation processes, such as RPS23 and RPL41.

## 5. Conclusion

We introduced a novel R package named EasyCircR, designed for comprehensive circRNA regulatory analysis, taking advantage of the post-transcriptional regulatory nature of circRNAs. Starting from RNA-seq FASTQ files of two distinct sample groups, our tool detects differentially expressed circRNAs between these conditions. It then identifies the miRNAs sponged by these circRNAs and determines their cascade target genes subject to post-transcriptional regulation. The analysis outcomes can be visualized, explored and downloaded through a Shiny app included in the package. We applied EasyCircR to analyze diffuse large B cell lymphoma cell lines treated with Bimiralisib, successfully validating the drug’s post-transcriptional regulatory impact. Additionally, we uncovered promising regulatory associations for potential further investigation. EasyCircR performance in terms of running time is closely tied to the usage of the CIRI-full tool for identifying and quantifying circRNAs. Moreover, in terms of detection power, the variability in sensitivity among circRNA detection tools is well-acknowledged [2]. To address this, EasyCircR will adopt a multi-tool approach within its pipeline to enhance the reliability of results through a consensus analysis. Additionally, our future plans include giving users the option to integrate several miRNA datasets into their analysis, which aims to provide even more realistic insights into regulatory interactions. Despite the limited understanding of circRNAs functions and the constraints of current analysis tools in this area, EasyCircR helps to make a step forward in understanding the regulatory role of circRNAs providing accurate and reliable associations.

## Notes

### Competing Interest Statement

The authors have declared no competing interest.

